# Effects of a temperate heatwave on diel rhythms of insect activity: a comparison across habitats

**DOI:** 10.1101/2025.01.08.631862

**Authors:** J.A. Carter, R.G. Davies, W.J. Nash, R.K.A. Morris

**Affiliations:** School of Biological Sciences, University of East Anglia, Norwich Research Park, Norwich, Norfolk, NR4 7TJ, UK; Earlham Institute, Norwich Research Park, Norwich, NR4 7UZ, UK; Scientific Associate, Diptera Section, The Natural History Museum, South Kensington, London, SW7 5BD, UK; Hoverfly Recording Scheme, 241 Commonside East, Mitcham, Surrey, CR4 1HB, UK

**Keywords:** Temperature, Heatwaves, Arthropods, Diel activity patterns, Microclimates, Refugia

## Abstract

1. The increasing frequency and intensity of heatwave events in temperate climates threatens to alter behavioural rhythms of ectothermic animals, such as insects. However, it is poorly understood how heatwaves affect daily activity patterns of insects, and whether shaded microclimates can moderate these responses.
2. We investigated impacts of a heatwave on the diel profile of insect activity, comparing effects across open, tree-covered and hedged habitats. Using yellow pan traps, insect activity was monitored from 07:00 to 19:00 on ten non-consecutive days, including two during a heatwave.
3. Insect counts exhibited a unimodal relationship with temperature.
4. During heatwave days, open habitat exhibited a significant (∼81.9%) reduction in counts compared to two ‘non-heatwave’ field-days, one before and one after the heatwave. Smaller, non-significant reductions were observed in the tree-covered (38.3%) and hedged (17.8%) habitats.
5. The diel activity profile on non-heatwave days approximated to a unimodal relationship, with GLMM-estimated counts peaking around 15:00; by contrast, heatwave days exhibited a bimodal profile, with predicted counts highest in the morning and evening.
6. Such heatwave-induced deformations of activity patterns, modelled as interactions between heatwave and time-of-day, were significant across all three habitat types.
7. The findings suggest that temperate heatwaves can markedly decrease insect activity levels, and that whilst shade-providing vegetational features may reduce this effect, diel patterns of activity are affected landscape-wide. As heatwaves become more frequent, preservation of trees and hedges in temperate landscapes is likely crucial to support resilience of insect activity and wider ecosystem functioning.

## INTRODUCTION

Climate change is an ever-growing issue for global biodiversity. There was a steady increase in global surface temperatures between 1951 and 2010, of which more than half was likely due to human emissions (IPCC, 2014); such warming has since continued (IPCC, 2023). Of concern is not only a rise in average temperatures, but also the growing threat of heatwaves - periods of elevated temperatures relative to normal conditions. Climate change has resulted in an increase in heatwave frequency and duration across Europe (IPCC, 2014, 2023) and in the UK specifically (Kendon *et al*., 2024), and these changes are expected to intensify with further warming (IPCC, 2023; Sanderson & Ford, 2016). It has therefore never been more important to understand the impacts of temperature rise and heatwaves on the functioning of the biosphere.

Ectothermic animals are particularly vulnerable to temperature fluctuations: unable to thermoregulate by direct physiological means, they instead rely on the temperature of their environment to maintain their body temperature within a suitable range for survival and metabolism (Mellanby, 1939). Insects are an especially vulnerable group, as they are generally small, with a large surface-area-to-volume ratio, and thus internal temperatures that are particularly dependent on temperature of their immediate environment (Clench, 1966; Kemp & Krockenberger, 2004). As postulated by Abram *et al*. (2016), effects of temperature on insects (and ectotherms in general) can be divided into (i) kinetic (‘bottom-up’) effects, whereby temperature constrains tissue-level physiological performance, amounting to impacts on organism-level fitness, and (ii) integrated (‘top-down’) effects, whereby insects use adaptations to regulate body temperature, thus mitigating unfavourable bottom-up effects. This model provides a causal pathway for the thermal sensitivity of insects.

Starting with the bottom-up effects: in an ectotherm, the relationship between body temperature and performance of physiological functions generally follows an asymmetric curve (Huey & Stevenson, 1979; Figure 1). Low temperature limits metabolism (therefore energy production) (Mellanby, 1939), whilst temperature above an insect’s ‘thermal optimum’ damages the endocrine system (Jankovic-Hladni *et al*., 1983; Rauschenbach, 1991), and accelerates water loss (Ahearn, 1970). This temperature-dependence of tissue-level physiology amounts to impacts at the level of the whole insect (Huey & Stevenson, 1979): below the thermal optimum, increasing temperature has been shown to facilitate locomotion (Berwaerts & Van Dyck, 2004; Forsman, 1999), and to accelerate feeding and growth rates (Kingsolver & Woods, 1997; Lee & Roh, 2010), whilst temperatures above the optimum can delay metamorphosis (Denlinger & Yocum, 1999) and reduce fertility (Proverbs & Newton, 1962). Overall, physiological functioning in both adult and larval insects can be constrained by temperatures below and above their optima - the latter being of great relevance in an era of climatic warming.

**Figure 1.**
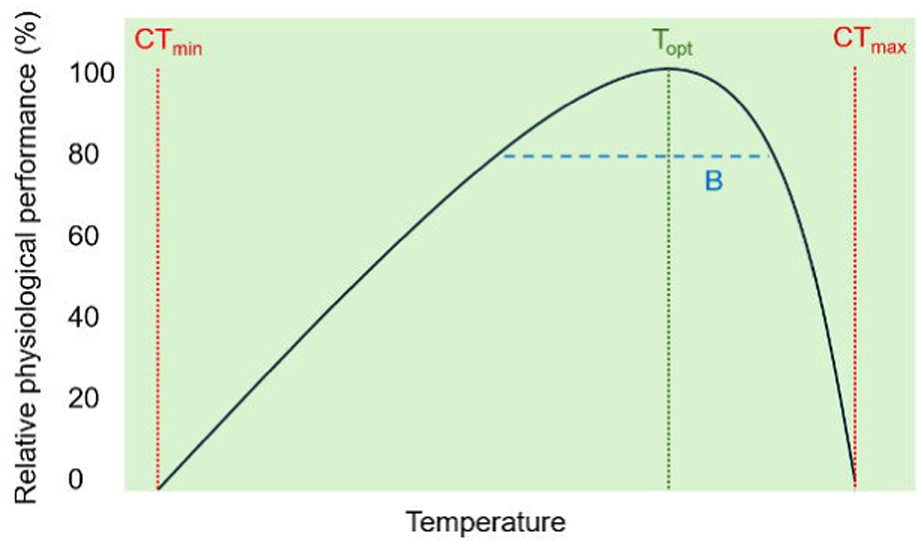
The thermal performance curve, which models the theoretical sensitivity of insect physiology to environmental temperature. The thermal optimum (T_opt_) is the temperature at which physiological performance reaches a maximum; the critical thermal maximum (CT_max_) and minimum (CT_min_) are the maximum and minimum temperatures at which any physiological functioning can occur; and the thermal performance breadth (B) is the temperature range across which an insect can perform ‘well’ - in this case, for example, at 80% or more of the maximum performance. Figure adapted from Huey & Stevenson (1979) and Huey *et al*. (2012).

Given such temperature-sensitivity of insect physiology, insects must necessarily utilise ‘top-down’ thermoregulatory mechanisms to remain within their thermal performance breadths (Figure 1). Ectotherms primarily rely on behavioural, over physiological, mechanisms to thermoregulate, adjusting their behaviour according to environmental temperature (Gunderson & Leal, 2015). This ‘behavioural plasticity’ has been observed repeatedly in insects in response to high temperatures. Heatwaves have been seen to cause marked reductions in bumblebee foraging time (Hemberger *et al*., 2023), butterfly activity (Hayes *et al*., 2024) and overall activity at ant colonies (Andrew *et al*., 2013), along with the retreat of aphids to gaps in the soil (Wiktelius, 1987), and of butterflies to shady refugia (Hayes *et al*., 2024). Overall, an understanding of ectotherm behavioural plasticity will be instrumental in understanding responses to global warming (Kearney *et al*., 2009; Sunday *et al*., 2014), and should be used to inform design and management of anthropogenic habitats.

Insect behavioural responses to heatwaves may be crucially influenced by habitat characteristics. Local- to landscape-scale heterogeneity in vegetation structure can create a diversity in thermal regimes that can be used by insects to thermoregulate (Sunday *et al*., 2014; Woods *et al*. 2015). Shaded areas under vegetation provide cooler refugia to which insects retreat in high temperatures (Clench, 1966; Hayes *et al*., 2024); such refugia could be crucial for ectotherm thermoregulation under climatic warming (Sunday *et al*., 2014; Woods *et al*., 2015). However, the way in which extreme events cause changes in insect activity across a habitat gradient is poorly understood; whether retreat to cooler refugia may allow insects to remain active during heatwaves, or whether these events may reduce activity landscape-wide. Heatwave-induced behavioural changes could interfere significantly both with insects’ population-level fitness and with their provision of ecosystem services – impacts that are likely to intensify if heatwave frequency, intensity and duration continue to increase. There is therefore a need to better understand how responses of insect activity to these events may be moderated by local habitat structure.

A key implication of the temperature-sensitivity of insect activity is that it translates into diel patterns, whereby activity changes with temperature through the day, with activity peaks and troughs varying based on latitude. At low latitudes, insects live close to their thermal optima (Deutsch *et al*., 2008). They should thus exhibit a bimodal pattern of activity through the day, with peaks in the morning and evening, and a trough around midday as insects avoid exposure to temperatures above their optima – a pattern observed in a Mediterranean climate by Herrera (1990). At higher latitudes, insects live further from, and less frequently exceed, their optima (Deutsch *et al*., 2008); thus, activity is limited primarily by low temperatures, and exhibits a unimodal trend, with one peak around midday, as seen in Alpine Norway by Totland (1994). However, even in a temperate climate, such as that of Britain, insects may be nearing their thermal optimum under global warming (Evans *et al*., 2019), and heatwave events could push insects above this optimum, provoking change in diel activity patterns. Temperate heatwaves have been observed to reduce insect activity (Hayes *et al*., 2024), but heatwave effects on rhythms of activity through the day are poorly understood. It is equally important to investigate how such impacts may vary across habitat types.

This study aims to examine the effect of a temperate heatwave event in Norfolk, UK, on diel patterns of insect activity, and how these patterns may vary between different structural habitat types. The following hypotheses are tested:

i. Insect activity should exhibit a unimodal relationship with temperature which corresponds with the thermal performance curve (Figure 1).
ii. Heatwaves should cause a reduction in total insect activity in open habitat, as insects seek refuge either by reducing activity or moving to shadier habitats.
iii. There will be an accompanying increase in total activity in the more shaded habitats (those with trees and hedges), as insects retreat to these refugia to resume activity.
iv. Heatwaves should change the general diel profile of activity from unimodal to bimodal, as high temperatures constrain activity around the middle of the day.
v. In shaded habitats specifically, this change will not occur, since temperature will not exceed the insects’ optima in these cooler microclimates.

We explore the implications of insect responses to the increasing threat of heatwaves. Investigating the extent to which shaded refugia can buffer these behavioural changes may inform habitat management on the potential need to integrate such habitats into open temperate landscapes to ensure ecological resilience. Ultimately, we find evidence that heatwave effects on total activity levels are reduced by presence of trees and hedges; however, we also find that heatwave conditions can transform diel rhythms of activity from unimodal to bimodal in profile, an effect which appears to occur regardless of local habitat structure.

## MATERIALS AND METHODS

### Data collection

The study site was a conservation area on the campus of the University of East Anglia, Norwich, Norfolk (Location: 52°36’36.0"N 52°37’12.0"E; see Figure 2). The site comprises a mosaic of habitats in a five-acre area, making it possible to walk easily between different habitats.

**Figure 2.**
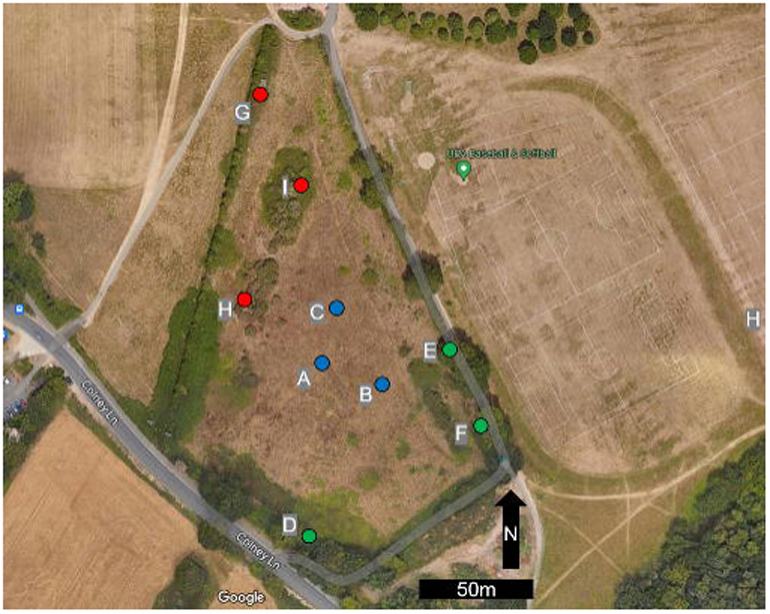
The study site (52°36’36.0"N 52°37’12.0"E). Circles show the sampled locations in the ‘open’ habitat (locations A, B, and C; shown in blue), the ‘trees’ habitat (D, E, F; green), and the ‘hedge’ habitat (G, H, I; red). Base map available at: https://www.google.com/maps

Following a preliminary visit to the site in June 2023, three structural ‘habitat types’ were coarsely mapped across it: open, tree-covered, and hedged areas (Table 1). Within each habitat type, three locations were then selected to be monitored throughout the study period, using a random co-ordinate generator. Measurements were later taken at each location to confirm that they conformed to their habitat-type definitions (Table 1). Distance to vegetation and vegetation height were measured using a tape-measure, whilst canopy cover for each site was estimated as the mean of four readings (north, south, east, west) taken using a spherical densiometer.

Insect activity was measured on ten non-consecutive days from July to September 2023 (Figure 3). Field days were selected opportunistically based on weather forecasts; an effort was made to capture days that spanned a wide variation in temperatures (including heatwave days), and to avoid rainfall, since this considerably reduces insect flight activity (Foster, 1974; Löhrl, 1976), potentially producing misleading results.

**Figure 3.**
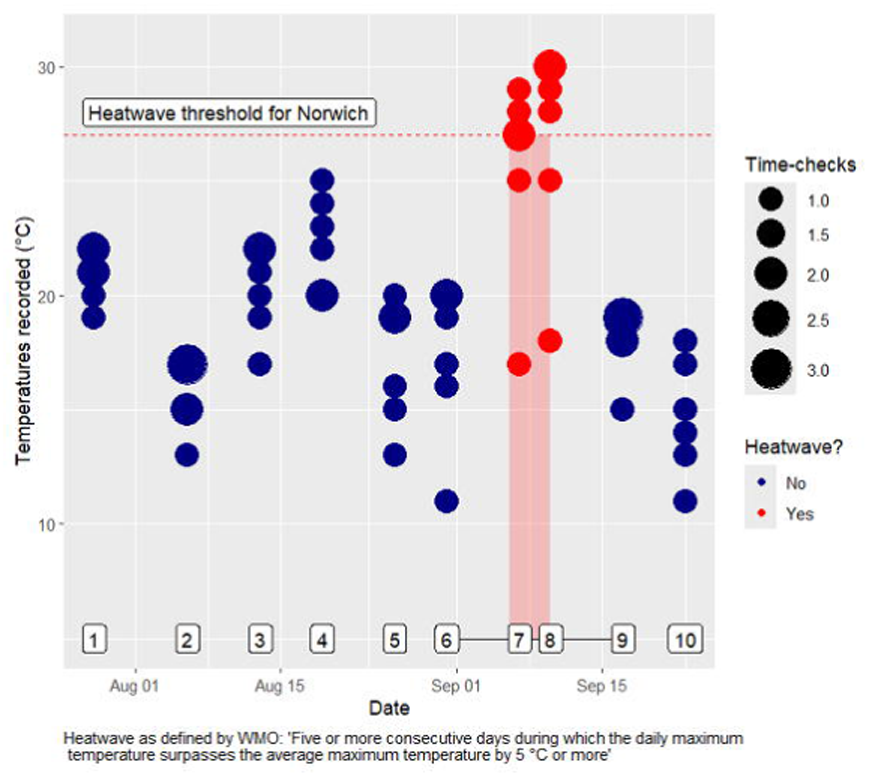
Temperatures observed at Norwich Weather Station on each of the ten field-days. For each of six bi-hourly time-checks (09:00-19:00), one reading was used (from halfway through the two-hour sampling window), so n=6 readings per day. The size of the dot reflects the number of occurrences of that temperature. The red dashed line shows the heatwave threshold temperature for Norwich. The sampled days are numbered 1-10 here, and are referred to as such in the text. The black solid line connecting days 6-9 shows the focal period for analyses of heatwave effects, whilst the red shaded rectangle shows the period over which the heatwave occurred (September 6^th^-10^th^). The dates of the ten sampled days were: July 28^th^; August 6^th^, 13^th^, 19^th^, 26^th^, 31^st^; September 7^th^, 10^th^, 17^th^, 23^rd^ (all 2023).

A heatwave, as defined by the World Meteorological Organisation, is “*five or more consecutive days during which the daily maximum temperature surpasses the average maximum temperature by 5°C or more”* (Rafferty, 2023). The average daily maximum for Norwich in summer is 22°C (Time and Date, 2024a); the threshold should thus be 27°C, which is also the heatwave threshold temperature currently used for Norfolk by the Meteorological Office (2022). The five days in the period 6^th^-10^th^ September (of which two were sampled) exceeded 27°C (Time and Date, 2024b; Figure 3); this period was thus considered a heatwave.

At each location selected for sampling, a pair of pan traps were placed to measure insect activity. The traps were plastic bowls filled with water and a small amount of scentless detergent. The latter reduces surface tension, facilitating submersion of invertebrates (Shrestha *et al*., 2019). The trap pair were placed one meter apart – sufficiently close that the vegetational environment surrounding each trap was near-identical. All bowls were painted on the inside with *Moon Glow* fluorescent yellow paint. Yellow pan traps have been shown to attract overall higher numbers of insects than other colours (Saunders & Luck, 2013; Vrdoljak & Samways, 2012). Furthermore, all traps were placed on the ground, as such an approach maximises counts and diversity of insects captured (Harris *et al*., 2017).

On each day, traps were deployed for twelve hours. They were laid and filled at 07:00, and subsequently checked at six two-hour intervals from 09:00 until 19:00. Each bi-hourly ‘time-check’ involved a 35-minute journey on foot between the nine sites. Sites were visited in the same order each time, such that all traps were left approximately two hours between checks. In each time-check, all invertebrate specimens in each trap were transferred to a labelled pot. Insects were then identified to order level, with use of a dichotomous key (Sanders & Provmsha, n.d.) where necessary, and counts of each order in each trap were recorded. Most identification was done in the field between time-checks, but order-ambiguous specimens were later identified in a laboratory.

To pair the observed insect counts with local weather, temperature data were taken from Norwich weather station (Time & Date, 2024b), which lies 7.4 kilometers from the study site. At the weather station, air temperature was recorded every 30 minutes. Each time-check was assigned the closest available temperature reading to halfway through the preceding two-hour sampling window (one hour before the time-check occurred). In addition, the duration of any rainfall that occurred within the two-hour window before each time-check was recorded *in-situ* using a stopwatch.

### Statistical analysis

Data visualisation and modelling were performed in R 4.3.3 software (R Core Team, 2021). The *tidyverse* package range (version 2.0.0) (Wickham *et al*., 2019) was used for data processing and visualisation, whilst package *glmmTMB* (1.1.8) (Brooks *et al*., 2017) was used for model-building.

Prior to modelling, the ten days were grouped into ‘heatwave’ and ‘non-heatwave’ treatments, according to the WMO definition mentioned previously. Days 7 and 8 (7^th^ and 10^th^ September) were placed in the heatwave treatment (Figure 3).

Generalised Linear Mixed Models (GLMMs) were used in all analyses (Supplementary materials, Table S1). The negative binomial distribution family was used, as all response variables were counts. Residuals of all models were confirmed, using function *simulateResiduals()* of package *DHARMa* (version 0.4.6) (Hartig, 2022) to adhere to the negative binomial distribution. In all models, three variables - Location, Day, and Hour nested within Day - were included as random effects (Supplementary materials, Table S1), to account for spatial and temporal non-independence. Function *r.squaredglmm()* of package *MuMIn* (version 1.48.4) (Bartoń, 2024) was used to estimate the proportion of variance in counts explained by each model’s fixed effects (the R^2^m value) and by its fixed and random effects combined (R^2^c). The lognormal forms of R^2^m and R^2^c were used, as appropriate for distributions with the logarithmic link function.

There is potential for spatial autocorrelation in model residuals to bias model results either via inflation of type I error rates and/or biasing of independent variable parameter estimates (Clifford *et al*., 1989; Cressie 1993). The residuals of all models were therefore tested, using Moran’s I, for spatial autocorrelation. Using function *correlog()* of package *ncf* (version 1.3-2) (Bjornstad, 2022), all 36 pairwise comparisons of the nine sites were grouped into distance-classes (of 40-meter increments), and Moran’s I and associated significance levels calculated for each distance-class. Focus lay on results for the 0-40m distance-class, for which P<0.05 would indicate a significant influence of short-distance spatial autocorrelation. No significant spatial autocorrelation in model residuals was observed in any of our models, allowing for reliable interpretation of independent variable effects (Diniz-Filho *et al*. 2003).

As a foundational basis for the investigation of heatwave effects, the effects of both temperature (at the mid-point of each two-hour sampling interval) and habitat type on insect counts were modelled first. This was done for total insect counts, and for counts of Diptera and Hymenoptera – the only two orders caught in adequate numbers for robust analysis (Supplementary materials, Table S2). The effect of temperature was tested by comparing linear and quadratic fit (lowest AIC is best), the latter with both a linear- and a squared-term simultaneously fitted as predictors. Preliminary examination of insect count distribution using package MASS (version 7.3-60.0.1) (Venables & Ripley, 2002) revealed that inclusion of data from day 10 (see Figure 3) caused insect counts to deviate from the negative binomial distribution (Supplementary materials, Figure S1); counts of Diptera on this day were markedly elevated compared to previous days (Supplementary materials, Figure S2), with a mean count per trap of 10.10 (the second-highest mean being 7.06 on day 6, and all other days exhibiting mean counts below 5). The increase was likely due to a seasonal ecological trigger beyond the scope of the study. To ensure good model fit, data from day 10 were excluded from the above analyses. In addition, to avoid misleading results, data from time-checks during which rainfall occurred were excluded. For all subsequent analyses, which examined heatwave effects, only a subset of the data was used. This included data from just four days - the two heatwave days (days 7 and 8), and the previous and subsequent, cooler field days (days 6 and 9, respectively) (see Figure 3). Use of such a seasonally proximal subset of days (ranging from August 31^st^ to September 23^rd^) minimises the influence of seasonality on insect counts, which could otherwise obscure behavioural patterns (Taylor, 1963). This data was used for three stages of modelling. Firstly, the overall effect of the heatwave on insect counts was modelled separately for each habitat type. Secondly, the relationship between time-of-day and insect counts was investigated, separately for heatwave and non-heatwave treatments. For each, the relationship was modelled as both linear and quadratic, and fits (AICs) of those two models compared. The final stage of the analysis was a culmination of heatwave, habitat, and time-of-day effects. Interactions were modelled between heatwave and both the linear- and squared-terms for time-of-day, to investigate whether heatwave presence significantly modified the shape (curvature) of the relationship between time-of-day and insect activity. This was modelled separately for all three habitat types.

## RESULTS

### Overview

Across the study period (n=1080 trap-checks), 6485 insects were trapped (average 6.00 specimens per trap-check). The most abundant insect orders were Diptera (4.67 specimens per trap-check), Hymenoptera (1.02), and Hemiptera (0.21). In the focal period for heatwave effects (days 6-9; n=432 trap-checks), 2106 insects were caught (4.88 per check). The most abundant orders were again Diptera (3.93 specimens per trap-check), Hymenoptera (0.76), and Hemiptera (0.10). (Supplementary materials, Table S2).

### The temperature-activity relationship and habitat effects

For all GLMMs modelling the effects of temperature and habitat type on counts (Table 2), inclusion of the squared-term for temperature alongside the linear term lowered the AIC (for total insects (a), from 4434.5 without the squared-term to 4410.7 with it; for Diptera (b), from 4001.4 to 3987.0; and for Hymenoptera (b), from 2255.8 to 2234.6). The squared-term was thus retained in the models to optimise fit.

Insect counts showed a unimodal, ‘concave-down’ (gradient decreasing) quadratic relationship with temperature (Figure 4a), shown by a positive estimate for the linear-term for temperature and a negative squared-term, with both terms significant (P<0.001) (Table 2a). Similar patterns were found specifically for Diptera (Figure 4b) and Hymenoptera (Figure 4c). In analyses of both Dipteran and Hymenopteran counts (Table 2b and c, respectively), temperature showed a positive coefficient and temperature^2^ showed a negative one, with both being significant (P<0.001).

**Figure 4.**
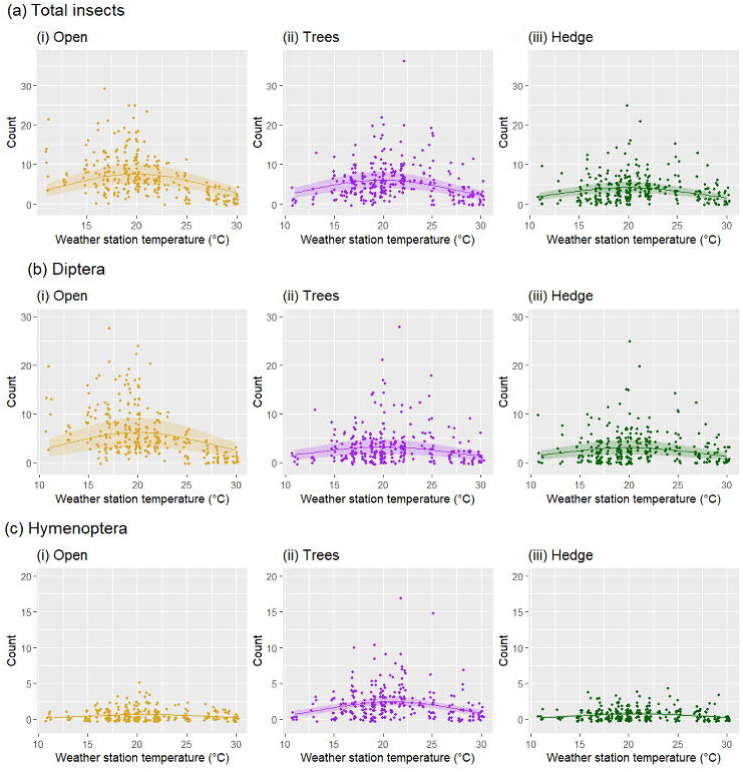
Relationships between air temperature (data from local weather station) and counts of insects caught in pan traps in a two-hour period (which the temperature reading was halfway through), for (a) total insects, (b) Diptera, and (c) Hymenoptera. For each taxon, n=864 trap-checks (288 per habitat type). Lines show estimates, and shaded bands show Wald 95% confidence intervals thereof, from corresponding Generalised Linear Mixed models for each taxon (Table 2), in which both temperature and habitat type are included as fixed effects; accordingly, data and model estimates are shown separately for (i) open, (ii) tree-covered, and (iii) hedged habitats. Models also include random effects (location, day, and hour nested within day). In all plots and their corresponding analyses, data from the final field-day (day 10) have been excluded, since their inclusion causes insect counts to violate the negative binomial distribution. In addition, data from two-hour windows where rainfall occurred were excluded. Note that the magnitude of the y-axis scales differs between the three taxa to enable clear visualisation of trends on their respective scales.

Total insect counts were highest in the open areas, being an estimated 19.9% lower in the tree-covered habitat and 44.8% lower in the hedged habitat (revealed by exponentiation of estimates from Table 2a); however, only the latter difference was significant (P<0.05). Dipteran counts, similarly, were highest in the open areas, with those in both tree-covered and hedged habitats exhibiting significant reductions (48.1% and 50.6%, respectively; both P<0.05) from the open-habitat intercept (Table 2b). For Hymenoptera, by contrast, counts were lowest in the open habitat, comparatively increased by 283% in the tree-covered habitat and 13.9% in the hedges; however, only the former difference was significant (Table 2c).

Habitat and temperature effects together explained an estimated 21.8% of the variance in total insect counts, 19.0% in Dipteran counts, and 35.3% in Hymenopteran counts (Table 2; see R^2^m values).

### Heatwave effects on activity levels

In all habitats, counts of insects caught decreased in the presence of a heatwave (Figure 5). Exponentiation of GLMM estimates in Table 3a reveals that, in the open areas (Figure 5a), count per trap decreased from 11.267 insects for the non-heatwave treatment (intercept) to 2.039 (18.1% of the non-heatwave count) for the heatwave treatment. This difference was significant (P<0.001) and explained 72.7% of the variance in counts (see R^2^m). In the tree-covered habitat (Figure 5b), there was a decrease from 4.417 insects per trap for the non-heatwave treatment to 2.989 (67.7%) during a heatwave (Table 3b). This difference was non-significant (P=0.107) and explained just 5.3% of variance in counts. In the hedged habitat (Figure 5c), there was a decrease from 2.749 insects per trap to 2.260 (82.2%) (Table 3c). This difference was also non-significant (P=0.285) and explained just 1% of variance in counts.

**Figure 5.**
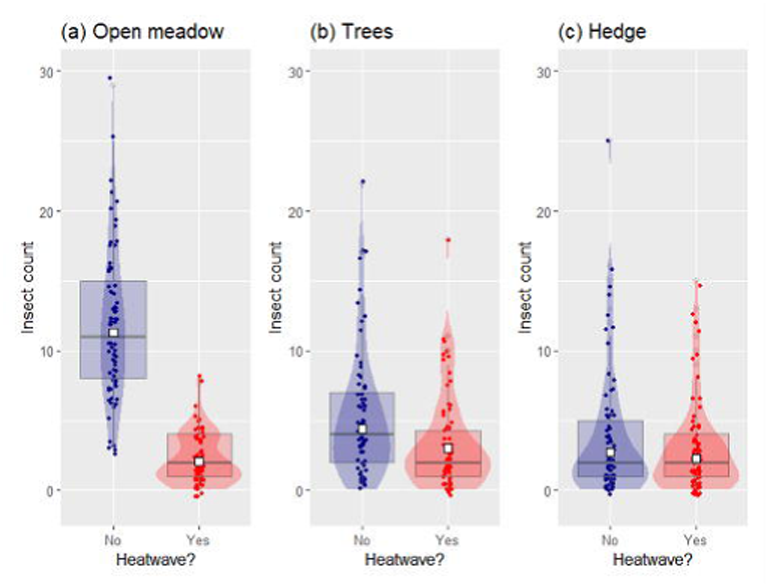
Counts of insects caught in pan traps during the heatwave (days 7 and 8) versus during two cooler days before and after (days 6 and 9), for (a) open habitat, (b) tree-covered habitat, and (c) hedged habitat (n=144 trap-checks per habitat, 72 per treatment). Raw data points show individual trap counts; boxplots show median and quartile range; violin plots visualise overall density distribution for each treatment; and white squares show estimates from corresponding Generalised Linear Mixed Models (Table 3), which include location, day, and hour nested within day as random effects. Whilst GLMM-estimated effects of the heatwave on insect counts were negative across all habitats, the decrease was significant (p<0.05) for the open habitat only.

### Diel temperature patterns: predicting effects on activity

On three of four days, temperature profiles were unimodal, with a peak at 14:00-16:00 (Figure 6). Day 9 showed a somewhat bimodal temperature profile.

**Figure 6.**
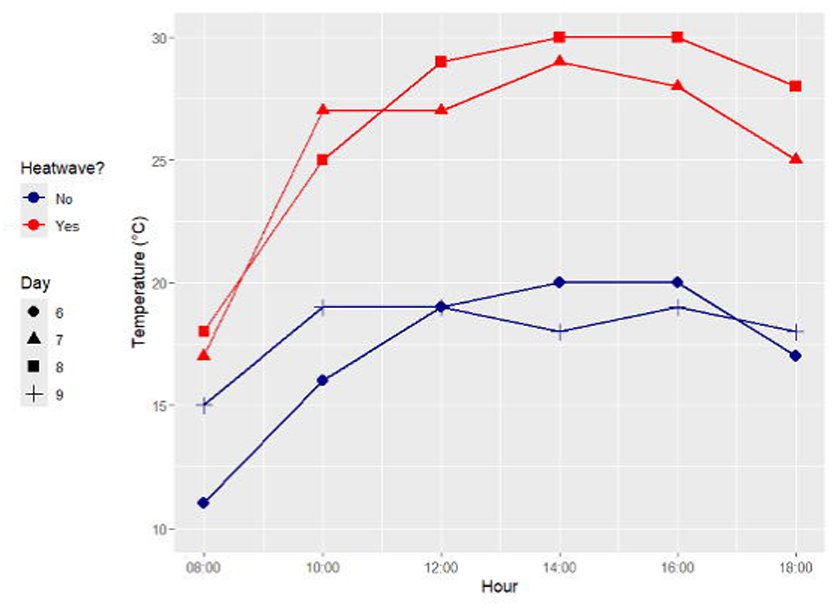
Observed changes in temperature across the four focal days (heatwave, days 7 and 8; non-heatwave, days 6 and 9), with weather-station temperature measurements taken halfway through each two-hour insect sampling window.

Using the observed diel temperature profiles (Figure 6), and model estimates of the temperature-activity relationship (Table 2a; Figure 4a), one can predict how insect activity should change through the day as temperature changes, and revisit earlier predictions on how heatwaves will affect such diel activity profiles. All temperatures observed on the non-heatwave days (Figure 6) were below the turning point of the temperature-activity curve (Figure 4a), with none exceeding 20°C; thus, one can retain the hypothesis that a unimodal profile of activity will occur on these days, as insect activity should never be limited by above-optimal temperatures, and should simply be highest when it is warmest (in this case, 15:00-17:00). By contrast, during the heatwave days, one can expect a bimodal pattern (as previously hypothesised), whereby activity declines from 9:00 onwards as temperatures exceed the optimum, and increases from 17:00-19:00 as temperatures decline toward the optimum (Figure 4a; Figure 6). Overall, the previously-generated hypothesis that heatwave conditions should change the diel distribution of insect activity from unimodal to bimodal can be retained for the next stage of the analysis.

Note that, as the theoretical thermal performance curve is asymmetric (Figure 1), but is modelled here as a quadratic curve (Figure 4a; Table 2a), the GLMM’s predictions should not be used to predict exact parameters for the temperature-activity relationship, but simply to verify and observe its general curvature.

### Diel activity patterns under heatwave and non-heatwave conditions

For both the non-heatwave (Table 4a) and heatwave (Table 4b) treatments, inclusion of a squared-term (hours after 7am^2^) in the modelling of time-of-day effects on counts lowered the AIC (for non-heatwave, from 1129.0 without the squared-term to 1128.1 with it; and for heatwave, from 905.5 to 901.8); thus, the squared-term, though non-significant in the non-heatwave model (P=0.064; Table 4a) was retained in both models to improve fit.

As predicted, a unimodal pattern of insect activity was observed on the non-heatwave days, and a bimodal pattern on the heatwave days (Figure 7).

**Figure 7.**
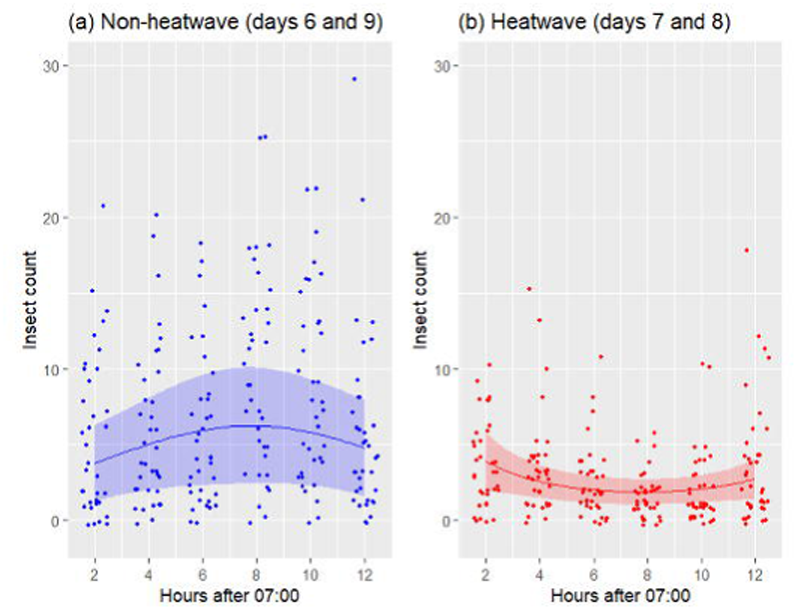
Counts of insects recorded in pan traps in six bi-hourly checks 09:00-19:00, (a) during the heatwave (days 7 and 8; n=216 trap-checks) and (b) during two cooler field days before and after (days 6 and 9; n=216 trap-checks). Lines show estimates, and shaded areas show Wald 95% confidence intervals, from the corresponding Generalised Linear Mixed Models (Table 4), which model quadratic effect of time-of-day on insect counts (separate models for heatwave and non-heatwave data). Location, day, and hour nested in day are included as random effects.

On the non-heatwave days, insect counts showed a concave-down (gradient decreasing) quadratic relationship with time-of-day, with GLMM-predicted count increasing to a peak around 8 hours after 07:00 (15:00) and declining thereafter (Figure 7a). Reflecting this concave-down effect, insect counts showed a positive relationship with the linear-term for time-of-day and a negative relationship with the squared-term (Table 4a). The effect of the linear term was significant (P=0.043), but that of the squared-term was not formally significant (P=0.064).

On the heatwave days, meanwhile, there was a concave-up (gradient increasing) quadratic relationship, whereby predicted count declined until around 8 hours after 07:00 (15:00) and increased thereafter (Figure 7b). Accordingly, there was a significant negative effect of linear time-of-day, and a significant positive effect of the squared-term, on insect counts (Table 4b).

Time-of-day effects overall explained a low proportion of the variance in insect counts (non-heatwave: 3.2%; heatwave: 7.5%) (Table 4).

### Habitat-specific heatwave effects on diel activity profiles

For all interaction terms shown in Table 5, AIC was increased by their removal from their respective models; thus, the explanatory power of these interactions more than compensates for the complexity they add to the models.

On the non-heatwave days, insect counts across all habitats exhibited a concave-down quadratic relationship with time-of-day as estimated by the GLMM (Figure 8). By contrast, and again consistently across habitats, model-predicted counts on the heatwave days exhibited a concave-up relationship with time-of-day (Figure 8). Across all three habitat types, heatwave presence interacted negatively with the linear-term for time-of-day, and positively with the squared-term, with significance (P<0.05) for all six interactions (Table 5). This suggests that the heatwave-induced inversion in the quadratic curvature of the diel activity profile, towards a concave-up relationship, was consistent across habitats.

**Figure 8.**
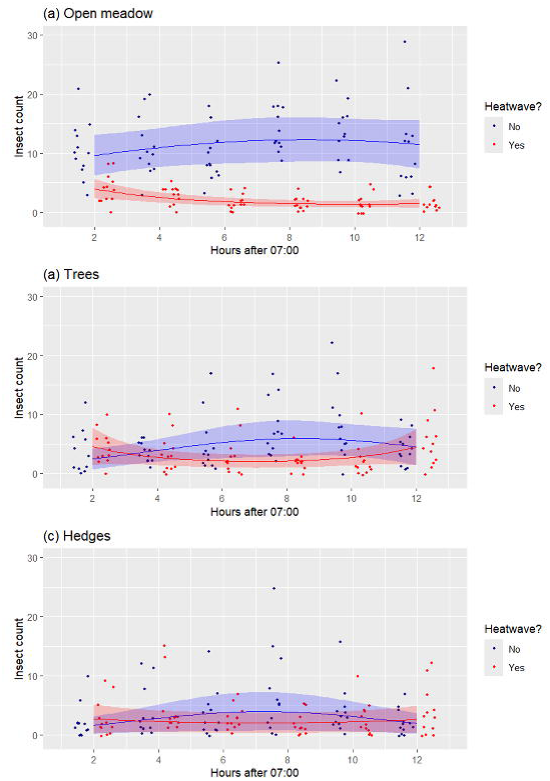
Counts of insects recorded in pan traps in six bi-hourly checks 09:00-19:00, during the heatwave (days 7 and 8; blue) versus during two cooler field days before and after (days 6 and 9; red), for (a) open habitat, (b) tree-covered habitat, and (c) hedges (n=144 trap-checks per habitat; for each one, n=72 for heatwave and 72 non-heatwave). Lines show estimates, and shaded areas show Wald 95% confidence intervals, from the corresponding Generalised Linear Mixed Models (Table 5), which model interactions between time-of-day (linear and quadratic terms) and heatwave presence, with location, day, and hour nested in day included as random effects.

## DISCUSSION

### Key findings

Activity of insects overall, and of orders Diptera and Hymenoptera, showed significant unimodal relationships with air temperature (Figure 4; Table 2). Examination of the heatwave versus non-heatwave comparison (Figure 5; Table 3) reveals significant temperature-dependence in the open habitat, where insect activity during the heatwave was markedly (81.9%) lower than on the two cooler days, with accompanying (albeit smaller and non-significant) reductions in tree-covered and hedged habitats. Such temperature-sensitivity also resulted in the diel rhythm of insect activity changing during the heatwave. Whereas counts approximated to a unimodal, ‘concave-down’ pattern on the cooler days, with a peak at 15:00, a bimodal, ‘concave-up’ profile was observed under heatwave conditions, whereby activity declined through the day, and subsequently increased in the evening as temperatures cooled (Figure 7; Table 4). Such interactions between heatwave presence and the gradient of the diel activity profile were consistent in direction and significance across all habitat types (Figure 8; Table 5).

### Temperature effects on activity levels

As hypothesised, there was a significant unimodal relationship observed between temperature and insect counts (Figure 4a; Table 2a), indicating constraints of above- and below-optimal temperatures on activity. This finding aligns with two key conceptual models. The first is the thermal performance curve (Huey & Stevenson, 1979; Figure 1): ectotherms’ physiological performance rises to a thermal optimum, and declines thereafter; and the second is that these ‘bottom-up’ temperature constraints provoke ‘top-down’ adaptations (Abram *et al*., 2016), whereby insects respond to environmental temperature (in this case, by reducing activity levels in above- and below-optimal temperatures) to maintain physiological functions. The unimodal temperature-activity relationship has been observed previously (Williams & Osman, 1960). The significant unimodal relationship observed consistently in this study across two major insect orders, Diptera (Figure 4b; Table 2b) and Hymenoptera (Figure 4c; Table 2c), supports a common influence of temperature in insect communities.

Specifically, there was strong evidence for constraints of extreme heat on activity (Figure 5). The substantial (81.9%) decrease in counts in open areas observed during the heatwave (Table 3a) supports the hypothesis that such events markedly reduce activity in these sun-exposed habitats. However, the hypothesised increase in activity in shaded (Trees and Hedge) habitats under the heatwave, during which insects were expected to retreat to these areas, was not observed, with decreases occurring in those habitats too (Tables 4b and c). These findings carry significant implications for the increasing threat of heatwaves under a warming climate. Whilst insect activity at temperate latitudes is overall predicted to increase with climate change, as temperatures rise closer to insects’ optima (Deutsch *et al*., 2008), the results above suggest that, if heatwaves are to become more common, then so too will periods of depressed activity. That decreases in activity for the tree-covered and hedged habitats were considerably smaller than for the open area (and non-significant) suggests that incorporation of these shaded refugia to a landscape should buffer overall heatwave impacts on activity. However, that the reduction in counts in open areas is not met with a corresponding increase in the shaded habitats suggests that such refugia may not offset the landscape-level loss of insect activity during heatwaves.

Such changes could profoundly impact insect populations and the ecosystems they support. Heatwave-induced reduction of insect activity may interfere with processes necessary for growth, survival or reproduction, threatening population-level fitness. For instance, Miler *et al*. (2020) showed that reduction in antlion foraging activity during an experimental heatwave also caused a reduction in body mass. Perhaps of even greater concern, however, are the wider ecological implications of insect activity reduction. Since the ability of insectivorous birds to catch arthropods is positively correlated with arthropod activity levels (Avery & Krebs, 1984), reductions in activity could reduce food availability for insectivores during heatwaves. Furthermore, a reduction in foraging by pollinators during heatwaves, as observed in bumblebees by Hemberger *et al*. (2023), could interfere with pollination services provided to plants. If heatwave frequency, intensity and duration continue to increase as predicted (IPCC, 2023; Sanderson & Ford, 2016), the above effects could increasingly interfere with the functioning of temperate ecosystems.

Furthermore, our findings warn of adverse ecological impacts potentially posed by the removal of hedges from agricultural landscapes across temperate Europe that has occurred in recent decades. Agricultural intensification during the 20^th^ century drove a substantial loss of hedgerows from European farmland (Poschlod & Braun-Reichert, 2017), with around half of Britain’s hedgerows removed (Barr & Parr, 1994). The resulting lack of shaded refugia to buffer effects of heatwaves on insect activity levels may threaten resilience at the population and ecosystem level. To mitigate this, we suggest that agricultural policymakers increase emphasis on retention and restoration of hedgerows in temperate European landscapes.

### Temperature effects on diel rhythms of activity

The diel profile of insect activity naturally varies across latitude, being generally unimodal at high latitudes, since insects do not typically experience temperatures above their optima (Totland, 1994), but bimodal in a lower-latitude Mediterranean climate, as high temperatures towards midday constrain activity (Herrera, 1990). The results of this study support the hypothesis that, by introducing such constraints of high temperature (Figure 6), a heatwave alters the diel profile of temperate insect activity (Figure 7; Table 4) from a unimodal form as seen on cool days, to a bimodal form fundamentally resembling that of a lower latitude. Furthermore, that such distortion in the diel profile of activity was consistent in direction and significance across all three habitat types (Figure 8; Table 5) would suggest that, contrary to what was hypothesised, temperatures during heatwaves may exceed insects’ optima and constrain their activity towards the middle of the day even under shaded refugia. Overall, it would appear that heatwave-induced change in the diel rhythm of activity is a landscape-wide effect that cannot be prevented by presence of shaded habitats.

Changes in diel patterns of insect activity under extreme temperatures may impact trophic interactions. If changes in activity times of insects are not matched by changes in foraging times in insectivores, insect and insectivore activity may become temporally mismatched. Such ‘trophic mismatch’ could exacerbate food shortages for insectivores during heatwave events. Insectivores can respond to gradual, seasonal changes in insects’ diel activity patterns by altering their foraging times through the year (Hódar *et al*., 1996), but whether they are able to respond this way to sudden emergence of heatwave conditions is unclear. The relative lack of studies of heatwave effects on trophic interactions makes it an important area for future investigations.

### Ambiguities and further research

Whilst this study provides valuable insight into temperature and heatwave effects on patterns in local insect activity, it does not explain the mechanisms underlying them. Changes in counts within a habitat could be due to movement of insects to or from that habitat, or simply changes in local activity levels. These are both potential ‘top-down’ temperature effects on insect behaviour observed previously under heatwave conditions (Hayes *et al*., 2024; Wiktelius, 1987). Lowered insect counts across *all* habitats during the heatwave suggest that such events do reduce overall levels of activity, rather than simply shifting it between habitats; however, the latter effect may also occur. Investigating the degree of insect movement that occurs across habitat gradients under heatwaves (for example, with use of flight interception traps, or by tracking individuals with remote sensing technologies) would help to disentangle potential explanatory mechanisms for the above findings.

The use of a ground-level sampling strategy limits the scope of the findings of this study. The tree-covered, and, to a lesser extent, hedged, habitats comprise taller vegetation, and therefore a greater three-dimensional volume of shade and resources, than open habitat. In these habitats, sampling at ground level accounts for just one layer (stratum) in a full vegetational column through which insects may be distributed. For both Diptera (Maguire *et al*., 2014) and Hymenoptera (Sobek *et al*., 2009), abundance in temperate woodlands can be greater at canopy level than ground level. This may explain why the heatwave-induced decrease in insect counts observed in the open habitat was not accompanied by the hypothesised increases under the tree-covered and hedged refugia: dissipation of any refuge-seeking insects through the full habitat volume may have resulted in a lack of any statistically significant change in counts detected at ground level. Overall, the conclusion that trees and hedges cannot balance the loss of insect activity in open habitats during heatwaves should be drawn with considerable caution, and further research, involving sampling at a range of vertical strata in the shaded habitats, will be necessary for a holistic understanding of the role of these refugia.

The specific effects of habitat are also unclear here. By dividing the site so coarsely into three ‘habitat types’, the study did not account fully for the vegetational characteristics of each site that may have influenced insects’ responses to heatwaves through shade or resource provision. In particular, the sites in the open habitat were casually observed to exhibit considerably higher densities of flowers in the proximity of the traps than those in the other habitats. Floral density positively affects site visitation of flower-visiting taxa such as bees and hoverflies (Hegland & Boeke, 2006; Hegland & Totland, 2005), and could explain, at least in part, why the baseline (non-heatwave) insect counts were highest in open areas (Figure 4). Meanwhile, since heat stress reduces nectar production in plants (Descamps *et al*., 2021), and thus insect visitation rates (Hemberger *et al*., 2023), the marked reduction in activity observed in the open, florally-dense areas during the heatwave may have been partially due to reduced local nectar provision, rather than lack of shade provision. Future research into variation in heatwave responses along distinct vegetational gradients (for example, canopy cover, plant diversity, and crucially floral density) would provide greater insight into the specific variables underpinning insect behaviour in extreme conditions.

Finally, future studies should examine heatwave responses of insects at greater taxonomic depth. Insects within an order are not physiologically uniform, and even closely-related taxa can differ in their responses to temperature and heatwaves (Martinez *et al*., 2023). For instance, hoverfly (Diptera:Syrphidae) species differ in their optimum temperatures for activity, and thus diel activity patterns, with larger species active at lower temperatures (therefore, earlier in the day) (Gilbert, 1985). Fine-scale variation in heatwave effects may exist between not only taxa, but also functional groups - for example, floral density is likely only to affect responses of flower-visiting taxa. Crucially, assessing effects of heatwaves on pollinator activity would allow prediction of the extent to which such events may impact agricultural productivity. Overall, our findings serve as foundations onto which further work should add insight with comparisons among taxonomic and functional groups.

## Conclusion

This study underscores the sensitivity of insect behaviour to extreme temperatures, and the importance of considering these ‘top-down’ temperature effects in models of ectotherms’ responses to climate change. Temperate heatwave conditions are shown here to markedly reduce levels and modify patterns of insect activity. Whilst we advise that creation and retention of shaded habitats in temperate landscapes should provide some relief against heatwave-induced activity reductions, this is unlikely to offset the reduction in open habitats, and diel rhythms are likely to be affected landscape-wide. Under ongoing climatic warming, these issues could become an ever-greater threat to the fitness of insect populations and the functioning of ecosystems to which their services are crucial.

Future research should investigate the specific top-down mechanisms driving the patterns observed here, disentangling inter-habitat movement from localised changes in insect activity. There is also a need to better understand how heatwave responses vary with distinct vegetational characteristics, giving particular consideration to floral density, and to account for the three-dimensional nature of shaded refugia by sampling at multiple strata. Finally, investigating these responses at family- or genus-level will provide deeper taxonomic insight. These developments would advance our understanding of the causal pathways underlying heatwave responses of insects, and how habitat-scale and landscape-scale conservation management could mitigate heatwave effects on global ecosystem functioning.

## ACKNOWLEDGEMENTS

The authors would like to thank the University of East Anglia for funding the purchase of materials required for field and lab work, as well as the university’s School of Environmental Sciences teaching lab team for providing laboratory space for specimen identification and sorting.

WN acknowledges support from the Biotechnology and Biological Sciences Research Council (BBSRC), part of UK Research and Innovation, Core Capability Grant BB/CCG2220/1 at the Earlham Institute and its constituent work packages (BBS/E/T/000PR9818 and BBS/E/T/000PR9819), and the Core Capability Grant BB/CCG1720/1 and the National Capability at the Earlham Institute BBS/E/T/000PR9816 (NC1—Supporting EI’s ISPs and the UK Community with Genomics and Single Cell Analysis), BBS/E/T/000PR9811 (NC4—Enabling and Advancing Life Scientists in data-driven research through Advanced Genomics and Computational Training), and BBS/E/T/000PR9814 (NC 3 - Development and deployment of versatile digital platforms for ‘omics-based data sharing and analysis).

## DATA AVAILABILITY

The dataset used in this manuscript, along with the R script used to analyse it, are available from Zenodo online repository (DOI: 10.5281/zenodo.13908996) (Carter *et al*., 2023). The dataset and code are not currently publicly visible, but can be viewed through this private link:

https://zenodo.org/records/13908996?token=eyJhbGciOiJIUzUxMiJ9.eyJpZCI6IjUyZjgzODgzLTdiYzItNDMyZS1hYTdkLTM1NTFhYTZhMjZlMSIsImRhdGEiOnt9LCJyYW5kb20iOiI0YmFlODE3MzQ4NWE5MTRhZWIyYjg3MTM5MzkwMGUwZSJ9.pL3yvakAae8J38dYAICFWPYmP4qaTR4-s7cVYvvMT9qfT0pWpnejqINYWB-t4sbtBKiYi4WBP77uBJl1xct9aw

*The full data citation is given in the reference list on the manuscript document*.

## USE OF ARTIFICAL INTELLIGENCE

ChatGPT-4 online generative AI software was used to assist in the development of various parts of the R code script used for data analysis. A list at the end of the script itself specifies which portions of code were developed with use of ChatGPT. No AI tools were used to develop any part of the manuscript itself.

## ETHICAL APPROVAL STATEMENT

The original project proposal received approval from the Animal Welfare & Ethical Review Board at the University of East Anglia.

## AUTHOR CONTRIBUTIONS

**Josh Carter:** Conceptualisation (equal); data curation (lead); formal analysis (lead); investigation (lead); methodology (equal); visualisation (lead); writing – original draft preparation (lead); writing – review and editing (equal). **Richard Davies:** Conceptualisation (equal); formal analysis (supporting); methodology (equal); supervision (equal); writing – review and editing (equal). **Will Nash:** Conceptualisation (equal); methodology (equal); supervision (equal); writing – review and editing (equal). **Roger Morris:** Conceptualisation (equal); methodology (equal); supervision (equal); writing – review and editing (equal).

*Note: table legends are included in the Word documents containing their respective tables*.

