## Supplementary Information for "Effects of a temperate heatwave on diel rhythms of insect activity: a comparison across habitats"

**Supporting information**

**Table S1.** An overview of all eleven Generalised Linear Mixed Models with negative binomial errors used in our analysis, and for each model, the variance calculated for each of the three fitted random effects: location of sample (9 levels), day of sample (10 levels), and hour nested within day (‘Hour:Day’; 6 levels of Hour per Day). All random effects were fitted only as random intercepts; no random slopes were fitted. The ‘Table relevant’ column indicates the table in the main text showing the fixed-effects output for each model. Fixed effects have been abbreviated as follows; ‘Habitat’ = habitat type; ‘Temp’ = temperature halfway through sampling window; ‘Hrs’ = hours after 7am. The ’Response’ column indicates the taxon of which counts are the model’s response variable. Shown in square brackets for each subset of the data used is the number of data-points (trap-checks).

| Table relevant | Fixed effects | Response | Data analysed [n] | Random effect | Variance |
| --- | --- | --- | --- | --- | --- |
| 2(a) | Temp; Temp^2^; Habitat | Insects | Days 1-9 (excl. rainfall-hours) [864] | Hour:Day | 0.022 |
|  |  |  |  | Day | 0.038 |
|  |  |  |  | Location | 0.064 |
| 2(b) | Temp; Temp^2^; Habitat | Diptera | Days 1-9 (excl. rainfall-hours) [864] | Hour:Day | 0.039 |
|  |  |  |  | Day | 0.073 |
|  |  |  |  | Location | 0.145 |
| 2(c) | Temp; Temp^2^; Habitat | Hymenoptera | Days 1-9 (excl. rainfall-hours) [864] | Hour:Day | <0.001 |
|  |  |  |  | Day | 0.063 |
|  |  |  |  | Location | 0.007 |
| 3(a) | Heatwave | Insects | Days 6-9; open [144] | Hour:Day | 0.062 |
|  |  |  |  | Day | 0.011 |
|  |  |  |  | Location | 0.027 |
| 3(b) | Heatwave | Insects | Days 6-9; trees [144] | Hour:Day | 0.265 |
|  |  |  |  | Day | <0.001 |
|  |  |  |  | Location | 0.060 |
| 3(c) | Heatwave | Insects | Days 6-9; hedge [144] | Hour:Day | 0.095 |
|  |  |  |  | Day | <0.001 |
|  |  |  |  | Location | 0.468 |
| 4(a) | Hrs; Hrs^2^ | Insects | Days 6&9 (non-heatwave) [216] | Hour:Day | 0.063 |
|  |  |  |  | Day | 0.039 |
|  |  |  |  | Location | 0.568 |
| 4(b) | Hrs; Hrs^2^ | Insects | Days 7&8 (heatwave) [216] | Hour:Day | 0.034 |
|  |  |  |  | Day | 0.032 |
|  |  |  |  | Location | 0.189 |
| 5(a) | Heatwave; Hrs; Hrs^2^; (Heatwave * Hrs);  (Heatwave * Hrs^2^) | Insects | Days 6-9; open [144] | Hour:Day | 0.011 |
|  |  |  |  | Day | 0.018 |
|  |  |  |  | Location | 0.027 |
| 5(b) | Heatwave; Hrs; Hrs^2^; (Heatwave * Hrs);  (Heatwave * Hrs^2^) | Insects | Days 6-9; trees [144] | Hour:Day | 0.159 |
|  |  |  |  | Day | 0.009 |
|  |  |  |  | Location | 0.061 |
| 5(c) | Heatwave; Hrs; Hrs^2^; (Heatwave * Hrs);  (Heatwave * Hrs^2^) | Insects | Days 6-9; hedge [144] | Hour:Day | 0.030 |
|  |  |  |  | Day | 0.001 |
|  |  |  |  | Location | 0.460 |

**Table S2.** Total counts of each insect order caught in the pan traps across all ten fieldwork days (n=1080 trap-checks) and across the focal period used for heatwave analyses, days 6-9 (n=432).

| Order | Total caught | |
| --- | --- | --- |
|  | **Days 1-10** (1080 checks) | **Days 6-9** (432 checks) |
| Diptera | 5048 | 1699 |
| Hymenoptera | 1103 | 330 |
| Hemiptera | 226 | 43 |
| Coleoptera | 70 | 20 |
| Orthoptera | 14 | 8 |
| Lepidoptera | 14 | 2 |
| Thysanoptera | 9 | 3 |
| Dermaptera | 1 | 1 |
| *Total insects* | *6485* | *2106* |

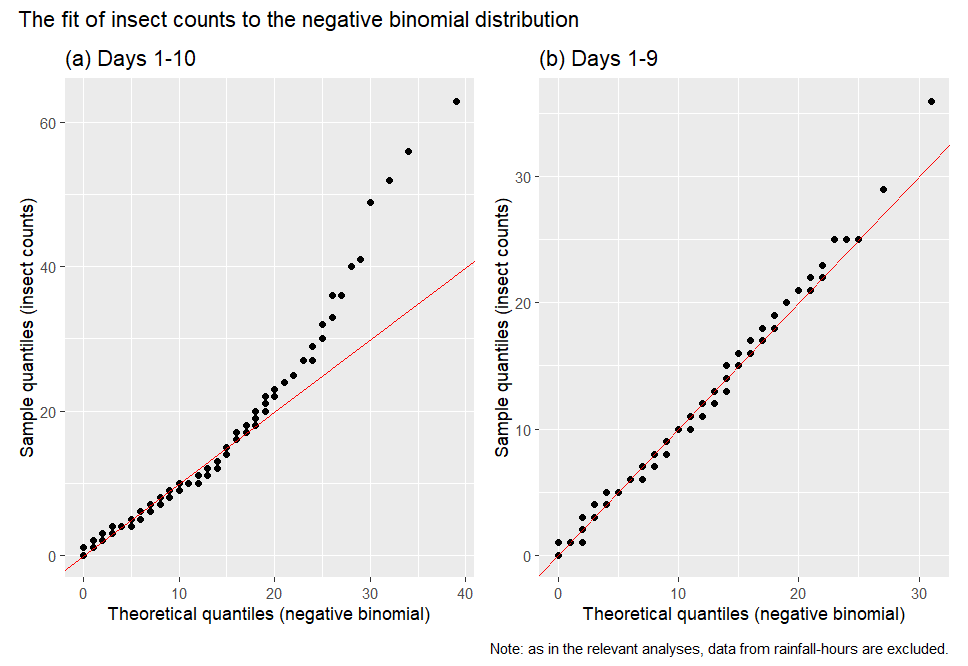

**Figure S1.** Q-Q plots comparing progression through quantiles of the distribution of observed insect counts to theoretical quantiles assumed by the negative binomial distribution, using (a) data from all ten field-days, and (b) data from all except the tenth field-day. Note that, as in the relevant analyses, data from rainfall-hours are not included here.

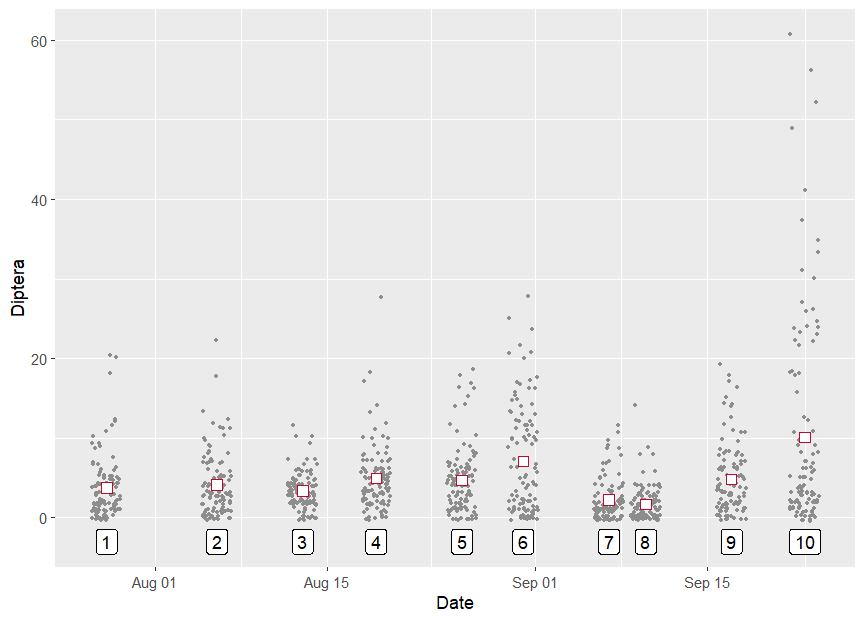

**Figure S2.** All per-trap counts of Diptera observed on the ten sampled days. For each day, the white square shows the mean count, and the label shows the number by which the day is referred to in the text.
